# Identification of cyclic hexapeptides natural products with inhibitory potency against *Mycobacterium tuberculosis*

**DOI:** 10.1101/279307

**Authors:** Sheo B. Singh, Joshua Odingo, Mai A. Bailey, Bjorn Sunde, Aaron Korkegian, Theresa O’Malley, Yulia Ovechkina, Thomas R. Ioerger, James C. Sacchettini, Katherine Young, David B. Olsen, Tanya Parish

## Abstract

A set of ∼500 purified natural product compounds was screened for inhibition against the human pathogen *Mycobacterium tuberculosis.* A series of cyclic hexapeptides with anti-tubercular activity was identified. Five analogs from a set of sixteen closely related compounds were active, with minimum inhibitory concentrations ranging from 2.3-8.9 μM. Eleven structural analogs had no significant activity (MIC > 20 μM) demonstrating structure activity relationship. Sequencing of resistant mutant isolates failed to identify changes accounting for the resistance phenotype.

---

Tuberculosis remains a global health problem, with an increasing number of new cases (10.4 million in 2015) and an increasing problem of drug resistance (1). New therapeutic agents which act against novel bacterial targets are needed to supplement and supplant the current drugs. Therefore there is great interest in finding new chemical matter that has activity against the causative agent, *Mycobacterium tuberculosis*. A large effort has been expended to examine small molecule libraries as starting point for drug discovery (2, 3). Less attention has been paid to natural products as starting points for a discovery program. We were interested in expanding chemical diversity to identify novel inhibitors which could be used either as starting points for a drug discovery program or as probes to identify novel targets.

A phenotypic screen to identify compounds with activity against *M. tuberculosis* using the MSD collection of purified natural products was initiated. A small set (∼500) of natural products for activity against *M. tuberculosis* H37Rv-LP (ATCC 25618) expressing a red fluorescent protein were tested (4-6). Compounds were tested at a single concentration of 20 μM. Growth was measured by fluorescence after 5 days of aerobic culture in Middlebrook 7H9 medium supplemented with 10% v/v OADC (oleic acid, albumin, D-glucose, catalase; Becton Dickinson) and 0.05 % w/v Tween 80 at 37°C in 384-well plates. Growth inhibition was calculated with reference to controls (DMSO only). Within this screen, hits were defined as having >30% inhibition of growth. Hits were confirmed by testing compounds in a dose response, using two readouts for growth – OD_590_ and fluorescence (Ex 560/Em 590). (4-6). Compounds were solubilized in DMSO, and a ten point, two-fold serial dilution prepared in DMSO. *M. tuberculosis* was cultured in Middlebrook 7H9 supplemented with 10% v/v OADC and 0.05 % w/v Tween 80. Cultures were inoculated at a final OD_590_ of 0.02 into 96-well plates containing compounds (final DMSO concentration of 2%). The starting concentration for compounds was 20 μM. Growth inhibition curves were plotted using the Levenberg-Marquardt algorithm and the concentration that resulted in 90% inhibition of growth was determined (MIC_90_). For active compounds, assays were conducted at least twice. Rifampicin was used as a control in each plate – for comparison the MIC of rifampicin was 0.0077 μM.

From these experiments, MSD-453 confirmed as a hit and further analogs based on similarity searches were ordered and tested. The initial hit, a cyclic hexapeptide, cyclo[N-(Me)Phe-Pro-Phe-N-(OH)Leu-Piz-Piz] (compound **1**) had good activity against *M. tuberculosis*, with an MIC_90_ of 2.3 μM (Table 1). Several natural congeners (**2-4)** and semi-synthetic and synthetic analogs (**5-16)** were available from the MSD sample library and were tested for activity against replicating *M. tuberculosis* under aerobic conditions in liquid medium. Structural identities and purity of the compounds were verified prior to anti-tubercular testing. Four other analogs (**2-5**) exhibited intermediate inhibitory potencies (MIC_90_ of 2.8 - 8.9 μM), whereas eleven compounds (**6-16**) showed no inhibitory activity (MIC_90_ > 20 μM).

**Table 1.**
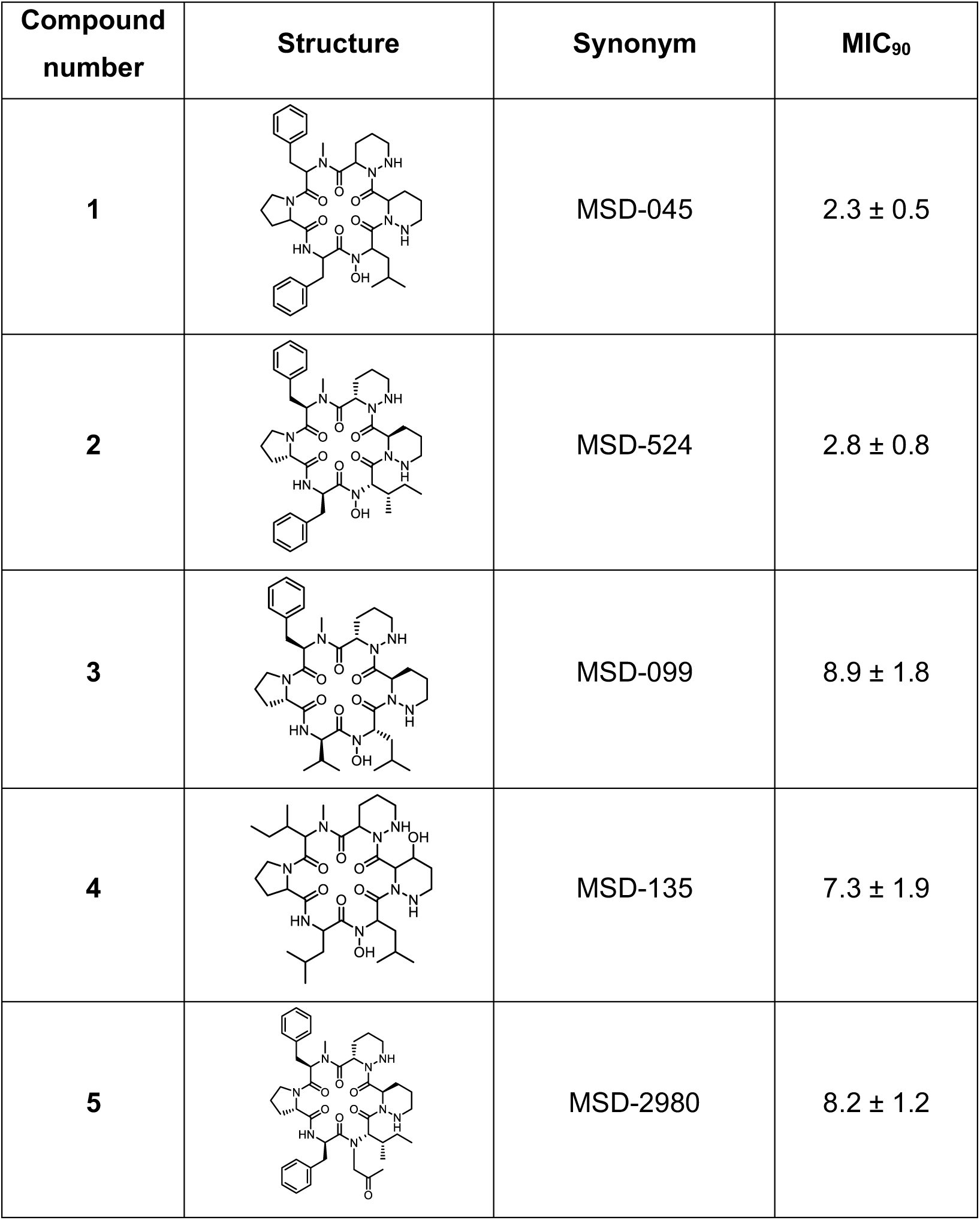
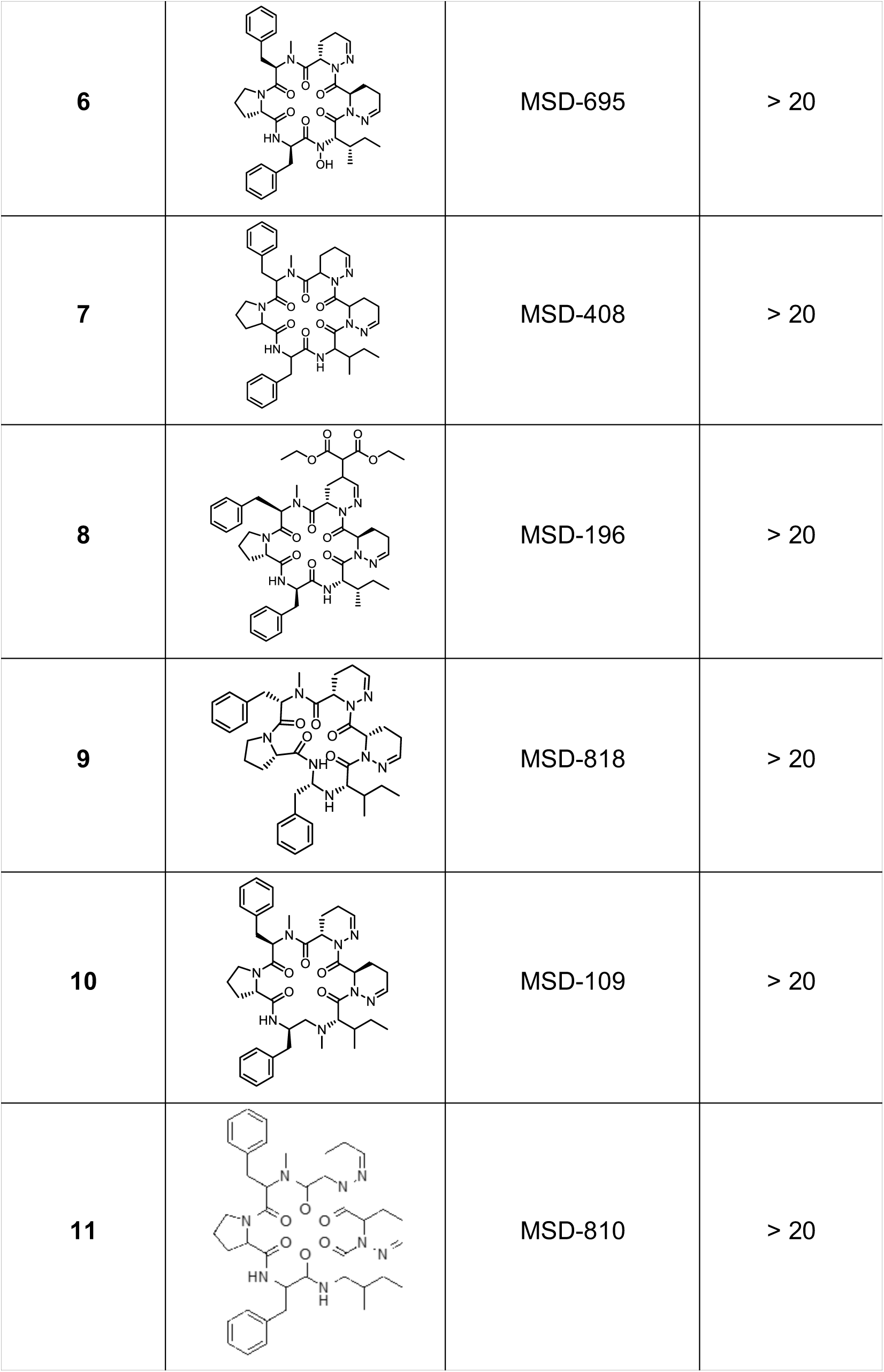
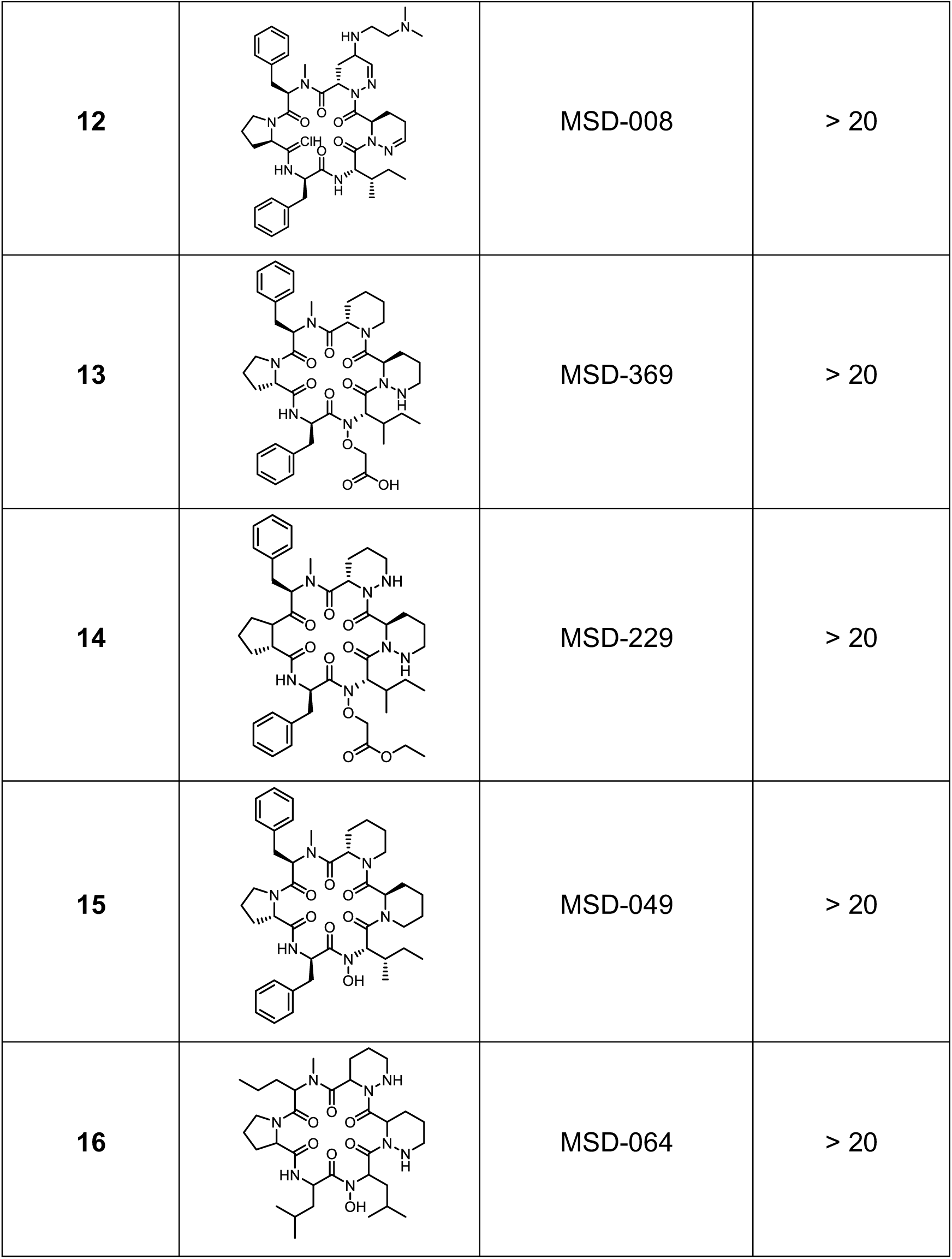
Activity of cyclic peptides against *M. tuberculosis*. MIC_90_ were determined against aerobically-cultured *M. tuberculosis* in liquid culture after 5 days of growth (5). Growth inhibition curves were plotted using the Levenberg-Marquardt algorithm and the concentration that resulted in 90% inhibition of growth was determined (MIC_90_). MIC_90_ were determined a minimum of two times for active compounds. Data are average ± standard deviation.

| Compound number | Structure | Synonym | MIC<sub>90</sub> |
| --- | --- | --- | --- |
| 1 | | MSD-045 | 2.3 ± 0.5 |
| 2 | | MSD-524 | 2.8 ± 0.8 |
| 3 | | MSD-099 | 8.9 ± 1.8 |
| 4 | | MSD-135 | 7.3 ± 1.9 |
| 5 | | MSD-2980 | 8.2 ± 1.2 |
| 6 |  | MSD-695 | > 20 |
| 7 |  | MSD-408 | > 20 |
| 8 |  | MSD-196 | > 20 |
| 9 |  | MSD-818 | > 20 |
| 10 |  | MSD-109 | > 20 |
| 11 |  | MSD-810 | > 20 |
| 12 |  | MSD-008 | > 20 |
| 13 |  | MSD-369 | > 20 |
| 14 |  | MSD-229 | > 20 |
| 15 |  | MSD-049 | > 20 |
| 16 |  | MSD-064 | > 20 |

Although the dataset of compounds tested are limited, some elements of the structure-activity relationship (SAR) (Figure 1) have been determined. Replacement of the N-hydroxyleucine with N-hydroxyisoleucine resulted in similar activity (**2**, MIC_90_ = 2.8 μM). Replacement of the phenylalanine with a valine reduced the activity slightly by 3-4-fold (**3**, MIC_90_= 8.9 μM). Similar reduced activity was seen after a double substitution of N-methylphenylalanine and phenylalanine with isoleucine and leucine (**4**, MIC_90_ = 7.3 μM). An N-alkyl-ketone in place of N-hydroxyleucine was tolerated, but reduced the biological activity by 3-fold (**5**, MIC_90_ = 8.2 μM), demonstrating that the N-hydroxyl group at that position is not critical. In contrast, the piperazic acids (Piz) were critical for activity of the molecule, since several tetrahydropiperazic (?Piz) acid derivatives (**6-12**) and the pipecolic acid (Pip) (**13** and **15**) analog resulted in loss of anti-tubercular activity. O-alkylation of the N-hydroxyleucine with an acetic acid or acetic acid ester abrogated activity (**13-14**). The combined replacement of N-methylphenylalanine with norvaline and phenylalanine with leucine also removed activity (**16**, MIC_90_ > 20 μM). Based on these analogs, we were able to determine some limited structure-activity relationship (Figure 1). Although the compounds were not as potent as rifampicin, they did have good activity, with a dynamic SAR ranging from low micromolar to inactive (>20 μM).

**Figure 1.**
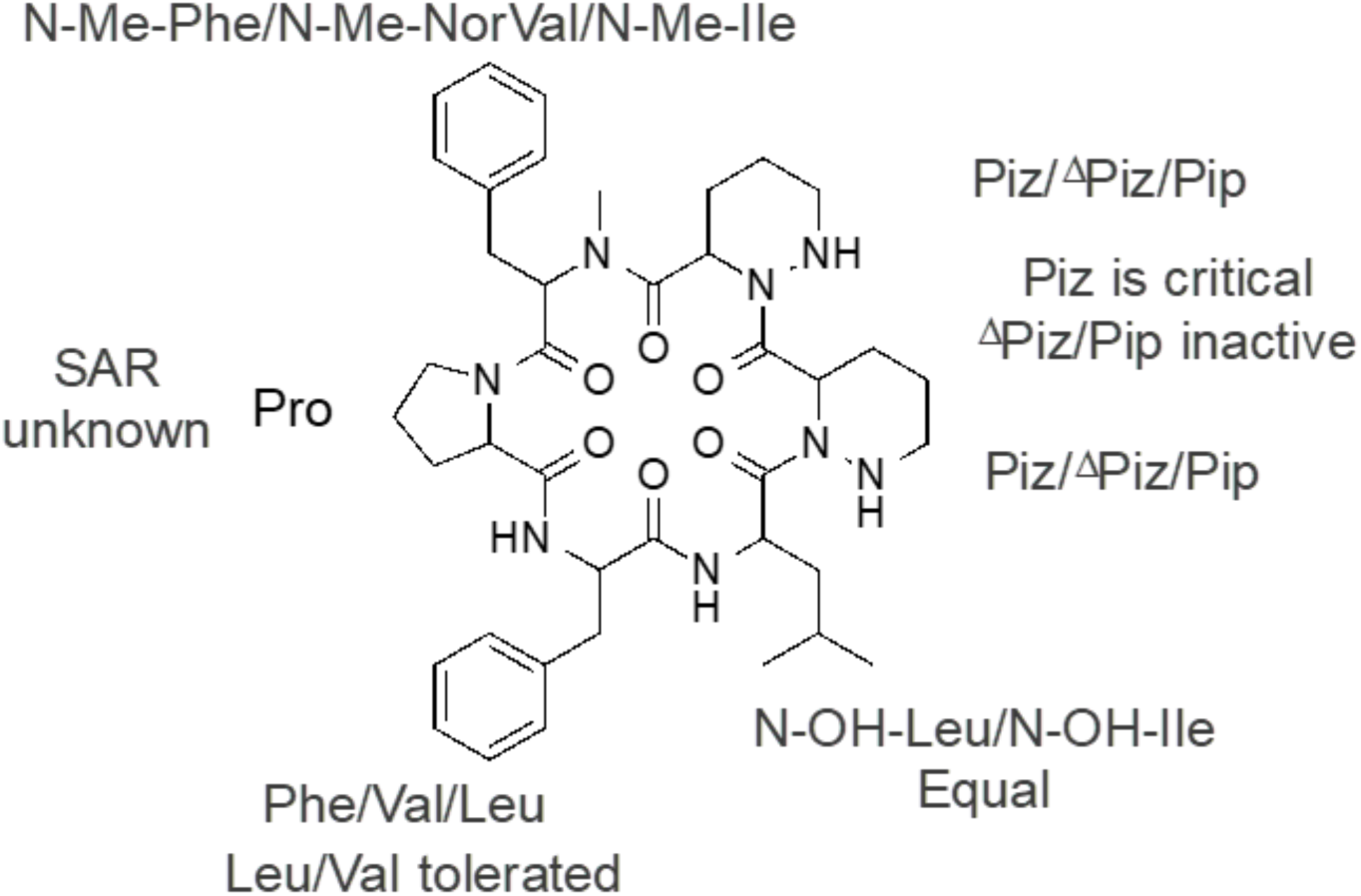
*M. tuberculosis* cyclic hexapeptide structure-activity relationship. Abbreviations: N-OH-Leu, N-hydroxyleucine; N-OH-Ile, N-hydroxyisoleucine; Phe, phenylalanine; Val, valine; N-Me-Phe, N-methylphenylalanine; Ile, isoleucine; Leu, leucine; Piz, piperazic acid; ΔPiz tetrahydropiperazic acid; Pip, pipecolic acid; N-Me-Nor Val, N-methyl-norvaline.

The original hit molecule, MSD-453, along with several congeners (10-13) and synthetic analogs (11-15), were originally identified as sub-micromolar to low nanomolar oxytocin receptor antagonists. *M. tuberculosis* does not have a homolog of this receptor; thus, this series appears to have a different target in the pathogen.

In an attempt to identify the *M. tuberculosis biochemical* target, resistant mutants to two of the compounds were isolated. The potency of each compound on Middlebrook 7H10 medium containing 10% v/v OADC supplement was measured. MIC_99_ values were determined by the serial proportion method and defined as the minimum concentration required to prevent 99% growth on solid medium (7). Under these incubation conditions, the MIC_99_ was 1.6 μM and 0.8 μM for MSD-045 and MSD-099 respectively (Table 2). Resistant mutants were isolated on Middlebrook 7H10 plus 10% v/v OADC at 5X MIC of each compound after 3-4 weeks growth (8). Colonies were tested for MIC_99_ on solid medium to confirm resistance (Table 2). In this longer term assay on solid media both compounds showed better potency than when tested for shorter time (5d) assay in liquid media (Table 1). Although nine confirmed resistant isolates were subjected to whole genome sequencing (9), no mutations were observed in 8 of the strains. Strain LP-0458686-RM1 had two mutations, one in *gap* (Rv1436) and one in Rv1543, but these were not observed in any of the other eight strains. Thus no genotypic explanation for the observed resistance was determined. In this case, the resistance phenotype could be due to epigenetic phenomena.

**Table 2.** *M. tuberculosis* strains resistant to hexapeptide inhibitor compounds. MIC_99_ were determined against *M. tuberculosis* resistant isolates on solid medium after 3-4 weeks. MIC_99_ was determined by the serial proportion method and defined as the minimum concentration required to prevent 99% growth (7).

| Strain | Compound | MIC<sub>99</sub> (μM) |
| --- | --- | --- |
| WT | MSD-045 | 1.6 |
| LP-0458690-RM1 | MSD-045 | 13 |
| LP-0458690-RM2 | MSD-045 | 25 |
| LP-0458690-RM3 | MSD-045 | 25 |
| LP-0458690-RM4 | MSD-045 | 25 |
| LP-0458690-RM5 | MSD-045 | 25 |
| LP-0458690-RM6 | MSD-045 | 25 |
| WT | MSD-099 | 0.8 |
| LP-0458686-RM1 | MSD-099 | 3.1 |
| LP-0458686-RM2 | MSD-099 | 3.1 |
| LP-0458686-RM3 | MSD-099 | 3.1 |

In conclusion, a series of cyclic natural product peptides with activity against *M. tuberculosis*, which could form the basis of further exploratory work including target identification, are disclosed. Future work to identify the intracellular target would assist in further development of this class of molecules.

## Acknowledgements

We thank James Ahn and Juliane Ollinger for technical assistance. This work was carried out under the auspices of the TB Drug Accelerator consortium.

## Funding

This work was supported in part by funding from the Bill and Melinda Gates Foundation under grant OPP1024038.

